# Integrated Collection of Stem Cell Bank data, a data portal for standardized stem cell information

**DOI:** 10.1101/2020.08.26.263830

**Authors:** Ying Chen, Kunie Sakurai, Sumihiro Maeda, Tohru Masui, Hideyuki Okano, Johannes Dewender, Stefanie Seltmann, Andreas Kurtz, Hiroshi Masuya, Yukio Nakamura, Michael Sheldon, Juliane Schneider, Glyn N. Stacey, Yulia Panina, Wataru Fujibuchi

## Abstract

The last decade has witnessed an extremely rapid increase in the number of newly established stem cell lines. However, due to the lack of a standardized format, data exchange among stem cell line resources has been challenging, and no system can search all stem cell lines across resources worldwide. To solve this problem, we have developed Integrated Collection of Stem Cell Bank data (ICSCB) (http://icscb.stemcellinformatics.org/), the largest database search portal for stem cell line information, based on the standardized data items and terms of the MIACARM framework. Currently, ICSCB can retrieve >16,000 cell lines from four major data resources in Europe, Japan, and the United States. ICSCB is automatically updated to provide the latest cell line information, and its integrative search helps users collect cell line information for over 1,000 diseases including many rare diseases worldwide, which has been a formidable task, thereby distinguishing itself from other database search portals.

## INTRODUCTION

Since the first report of human induced pluripotent stem cells (iPSCs) (Takahashi et al., 2007), there has been a rapid increase in the number of iPSC lines and related information worldwide **(Table 1)**. This remarkable growth has not only accelerated studies of regenerative medicine but also provided opportunities to understand such pragmatic issues as the quality of pluripotent stem cells (Nishizawa et al., 2016) and the disease mechanisms (Sasaki et al., 2016). Stem cell banks and registries are expected to provide necessary data of individual stem cell lines. However, the exchange of data among different institutions is not a trivial matter, and the scientific reproducibility of the stem cells, particularly iPSCs generated by different methods, depending on available information is problematic for both basic studies and clinical applications (Yaffe et al., 2016; Isasi and Knoppers, 2011; Thirumala et al., 2009). Moreover, as technologies for the characterization of cell lines continue to advance, the addition of new quality standards as necessary data items has complicated and diversified data formats among different stem cell banks and registries (Hug, 2009; Knoppers and Isasi, 2010). As an attempt to solve these problems, we previously reported MIACARM (Minimum Information About a Cellular Assay for Regenerative Medicine) guidelines in 2016 (Sakurai et al., 2016), which proposed the utilization of standardized data items and formats for all stem cell lines in regenerative medicine. Presently, MIACARM contains 260 items covering such areas as stem cell production and materials (e.g., donor information, source cell information, and cell culture medium and substrate information), cell banking processes, cell characterization, sterility testing, and even ethical concerns. Later, a standardized nomenclature for pluripotent stem cells was introduced in 2018 with unification of cell line codification and minimization of information loss and confusion regarding cell lines as goals (Kurtz et al., 2018). Nevertheless, with the growing number of registered cell lines, existing data deposition formats have made it increasingly harder for not only data depositors but also users to seek and obtain cell lines collected under different projects, disease states, and privacy issues (Godard et al., 2003; Winickoff et al., 2009).

**Table 1.** **Stem cell banks and registries worldwide (as of December 6, 2020)**

| B/R* | Stem cell bank or registry | Country | Website | Number of cell lines | If it has data included in ICSCB's registries (○: included, △: planned) |  |  |
| --- | --- | --- | --- | --- | --- | --- | --- |
|  |  |  |  |  | SKIP | eagle-i | hPSCreg |
| B | BLCB | Spain | https://p-cmrc.cat/ | 176 |  |  |  |
| R | hPSCreg | Germany | https://hpscereg.eu/ | 3360 | ○ |  |  |
| B/R | HipSci | United Kingdom | http://www.hipsci.org/ | 3720 |  |  | ○ |
| B | EBiSC | Germany | https://ebisc.org/ | 897 |  |  | ○ |
| B/R | CIRM / FUJIFILM | United States | https://fujifilmcdi.com/the-cirm-ipsc-bank/ | 1554 |  |  | △ |
| B | Harvard Stem Cell Institute | United States | http://stemcelldistribution.harvard.edu/ | 41 |  |  |  |
| B | NYSCF | United States | https://nyscf.org/ | 111 |  | ○ |  |
| B | NINDS Human Cell and Data Repository | United States | https://bioq.nindsgenetics.org/ | 162 |  |  |  |
| B | WiCell Research Institute | United States | https://www.wicell.org/ | 1519 |  | ○ | △ |
| B/R | eagle-i | United States | https://www.eagle-i.net/ | 2415 | ○ |  |  |
| B/R | RIKEN BRC | Japan | https://www.brc.riken.jp/en/ | 4102 | ○ |  |  |
| R | SKIP | Japan | https://skip.stemcellinformatics.org/ | 5770 |  |  |  |
| B | JCRB | Japan | https://cellbank.nibiohn.go.jp/ | 31 | ○ |  |  |
| B | Taiwan Human Disease iPSC Service Consortium | Taiwan | https://catalog.bcrc.firdi.org.tw/ | 102 | ○ |  |  |
| B | National Stem Cell Bank of Korea | Korea | http://kscr.nih.go.kr/ | 147 |  |  |  |

\* B, bank; R, registry

In this paper, as our next step towards the unification and utilization of stem cell line data in the world, we report our new database portal, Integrated Collection of Stem Cell Bank data (ICSCB), which was designed using MIACARM guideline items and formats, to serve an entrance ‘port’ to individual data repositories. The main objectives of ICSCB are i) to establish an integrated stem cell database portal that can cover the majority of stem cell resources in the world, and ii) to offer users minimum but efficient access to information on stem cell lines based on MIACARM guidelines. Currently, ICSCB provides data of more than 15,000 stem cell lines registered in four major stem cell line databases: hPSCreg (Seltmann et al., 2015), SKIP (Kim et al., 2017), RIKEN BRC (Kobayashi et al., 2016), and eagle-i (Vasilevsky et al., 2012). ICSCB has a user-friendly search engine for stem cell lines and can be accessed directly at http://icscb.stemcellinformatics.org/, or as a slim version by removing cell line redundancy as much as possible through the SHOGoiN (Human Omics Database for the Generation of iPS and Normal Cells) homepage at http://shogoin.stemcellinformatics.org/.

## RESULTS AND DISCUSSION

### Web interface

ICSCB was designed for researchers searching for available cell lines to conduct various studies, such as regenerative medicine and disease analysis. Covering as many diverse cell lines as possible was the first priority when deciding which resources to include in ICSCB. Sharing cell line information between different stem cell banks and registries has been problematic due to different cell naming methods, different policies on cell assessment in different registries, unclear data sources, and so on. ICSCB is a collection of cell lines from four major and reliable cell line data resources based in Europe, Japan, and the United States. ICSCB updating is regularly performed for new SKIP and eagle-i stem cell line data as well as automatically performed for hPSCreg and RIKEN BRC data in a synchronized manner. Users can retrieve all related stem cell line information by using a free text search. Detailed information of a specific cell line can be accessed by clicking on the stem cell ID, which is linked to the information page in the original resources (**Figure 1**). The results can be further filtered according to users’ requests. There may be several records for the same cell line if the cell line is included in multiple data resources. To provide users as much information as possible, the results page is designed to show cell lines with matching cell names as well as similar descriptions.

**Fig 1:**
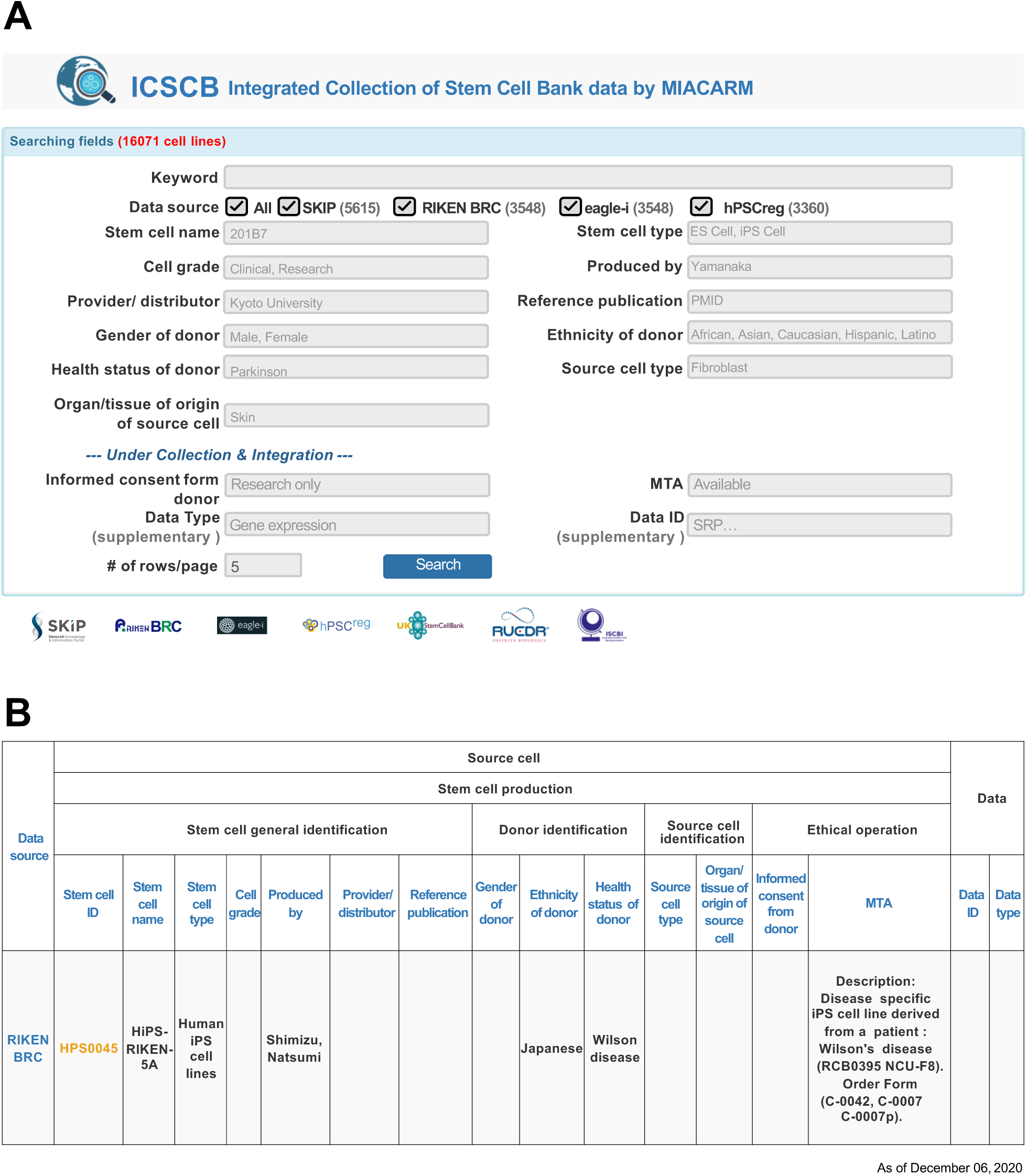
Web interface of ICSCB. (A) The ICSCB search page. Any keyword related to cell lines (including cell line name, disease name, gender, and so on) can be used to perform an instant search. (B) The ICSCB results page. Matched or partially matched cell lines are listed according to MIACARM terms. To check the details of the cell lines, the user can click on the stem cell ID, which is linked to the original source of cell line information.

### Data coverage

ICSCB covers more data than any other stem cell line repository available. The integration of all major data resources allows us to check the current state of stem cell research in the world **(Figure 2)**. Although we recognize redundancies in the data, according to our statistics, the number of iPSC lines constitutes more than 80% of all cell lines and the ratio of healthy to diseased donors is approximately 3 to 2 **(Figure 2A,B)**. The total number of countries from which cell lines can be retrieved is 39 (as of December 6, 2020), of which the top 9 countries identified in SKIP and hPSCreg are (in descending order) the United Kingdom, United States, Japan, Germany, China, Spain, Sweden, Denmark, and Taiwan **(Figure 2C)**. In addition, as the recent number of iPSC lines generated from patient donors is growing, ICSCB supports disease-oriented search to help users find all disease-related stem cell lines by using disease names. The distribution of disease and disorder types is shown in **Figure 2D**.

**Fig 2:**
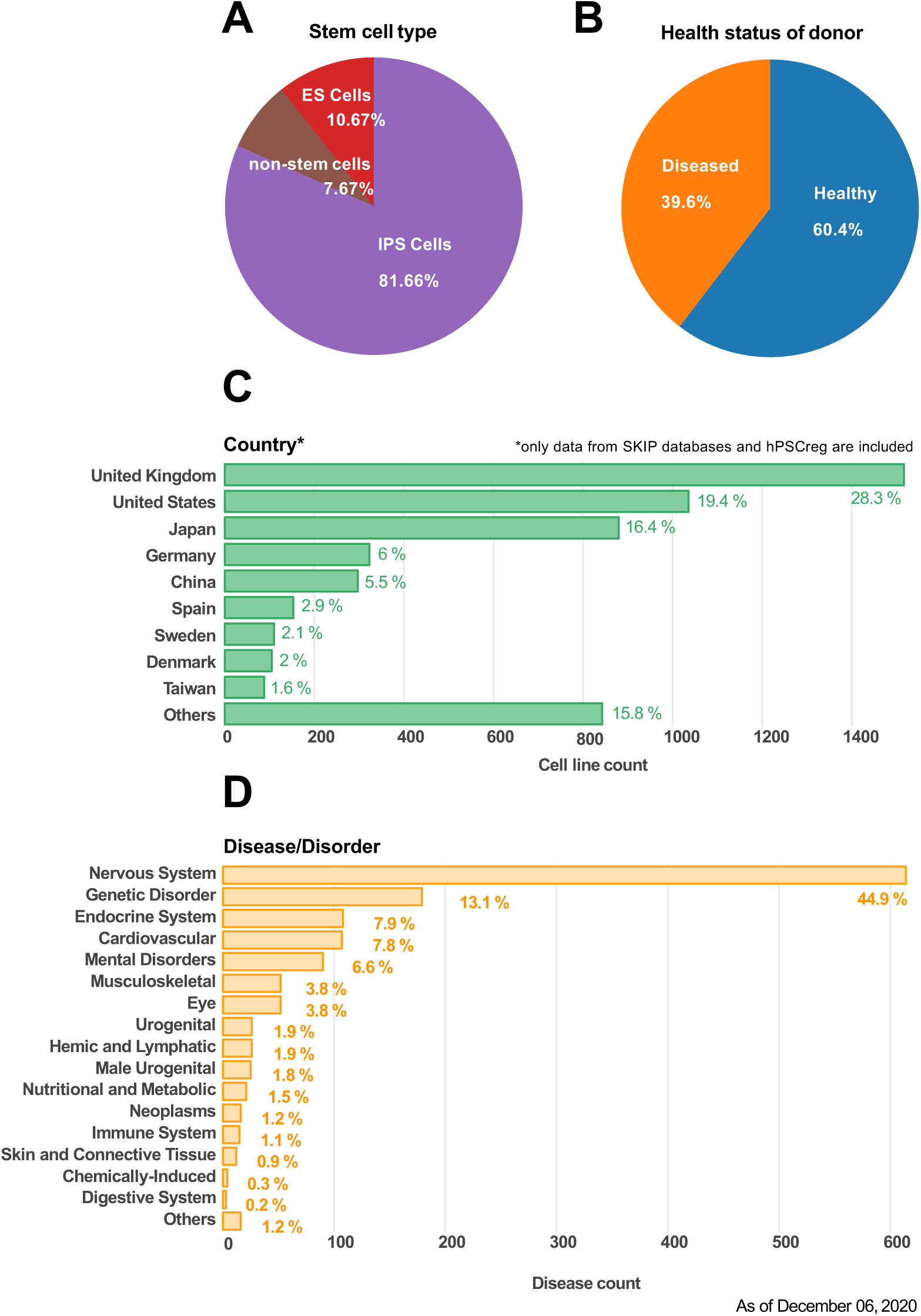
Details of cell lines collected by ICSCB. Cell line information is categorized as (A) stem cell type, (B) health/disease status of donor, (C) country that established the cell lines, and (D) disease category.

### Easy search interface on SHOGoiN homepage

ICSCB also has a quick and easy search module on the SHOGoiN homepage (https://stemcellinformatics.org/). SHOGoiN is a repository for accumulating and integrating diverse human cell information to support a wide range of research using cell-related data. The database consists of several modules that store cell lineage maps, transcriptomes, methylomes, cell conversions, cell type markers, and cell images with morphology data curated from public as well as contracted resources based on sophisticated cell taxonomy. Collaboration between ICSCB and SHOGoiN makes it possible for users to directly use free text searches for stem cell line data on the SHOGoiN homepage. The ICSCB easy search module in SHOGoiN supports a simplified ICSCB search with keywords, and the advanced search is designed to redirect users to the ICSCB homepage with full functions. Results from the SHOGoiN homepage share the same structure with the ICSCB homepage.

### Concluding remarks and future plan

So far, the registration and submission of newly established cell lines have been complicated by the lack of standardized data formats. Most data registries are currently limited by respective domestic policies and have adopted their information structures and validation processes independently (Andrews et al., 2015; Zarzeczny et al., 2009). The lack of standardized data formats has caused problems for researchers who must usually search several websites to find the stem cell lines they are looking for (Wells et al., 2013). In the present work, we developed ICSCB, an integrated data distribution system that provides stem cell line information from major stem cell banks and registries all over the world. ICSCB adopts a standardized information format based on the “Source Cell” module of MIACARM to integrate different data resources while keeping important information.

ICSCB has several limitations; for instance, there exist cell lines having limited donor information and/or incomplete information, as well as replicates of the same line with different names. In addition, ICSCB contains only a few clinical-grade cells due to strict requirements for exhaustive as well as expensive quality checks and haplotype compatibility for clinical-grade lines. Currently, when searching for stem cell lines for a specific disease, users have limited access to a distinct aspect of research for the diseases registered in individual repositories. Indeed, our next step would be to establish an integrated and refined collection of research on stem cell lines in order to understand the possible causes and mechanisms of complex diseases on the basis of genetic backgrounds and environmental effects in terms of molecular pathways during the developmental process. We may need a new project based on new funding to establish such a global collaboration.

ICSCB has several issues that merit improvement. Firstly, we plan to assign unique accession codes to all cell line entries by utilizing standardized nomenclature for pluripotent stem cells (Kurtz et al., 2018) in order to remove redundant cell line data. Secondly, as the cost of experimental technologies, such as genome/RNA-seq or teratoma assay, to characterize stem cell lines decreases and it becomes easy to obtain various types of profiles on them, we may be able to define a standard profile set for complete data format to render comparisons of cell lines more efficient. Thirdly, once the RNA-seq or genome mutation data are collected, it will be possible to perform statistical analysis such as PCA or other refined bioinformatics methods to mathematically map individual cell lines to a global stem cell feature space, such as differentiation propensity, carcinogenic potential, immune response, and so on. We expect ICSCB to further evolve, thereby providing users better accessibility to relevant stem cell lines.

In the future, in order to respond to the rapid growth in the number of stem cell lines, we will include more data resources in ICSCB, including the Taiwan Human Disease iPSC Service Consortium and other recently developed stem cell banks, to make ICSCB more resource-abundant and usable. We also plan to add a detailed quality check to help users find stem cell lines of high quality. As the largest stem cell line information resource, we will support stem cell communities by improving the quality and increasing the scale of our database.

## EXPERIMENTAL PROCEDURES

### Data resources

ICSCB resources were selected from existing major stem cell registries that collect cell line information in Europe, Japan, and the United States, and stem cell banks that provide cell lines with information of the attributes. We checked the number of registered cell lines and the criteria for registration in these registries and banks to decide to what extent their cell line data can fulfill MIACARM guidelines for inclusion in ICSCB. Considering the size, accessibility, and diversity of the different databases, three stem cell registries and one stem cell bank were included: (1) SKIP (5,615 cell lines), (2) hPSCreg (3,360 cell lines), (3) RIKEN BRC (3,548 cell lines), and (4) eagle-i (3,548 cell lines) (as of December 6, 2020). These data resources were selected because they had the highest number of registered cell lines and large diversity, which would provide a good regional balance of cell sources to reduce redundancies in cell line entries. RIKEN BRC basically collected cell lines from Japanese institutions, SKIP contained data mostly from other Japanese and Asian institutions, hPSCreg collected data mainly from European institutions, and eagle-i collected data mostly from the United States. Details of the data sources are listed in **Figure 3**.

**Fig 3:**
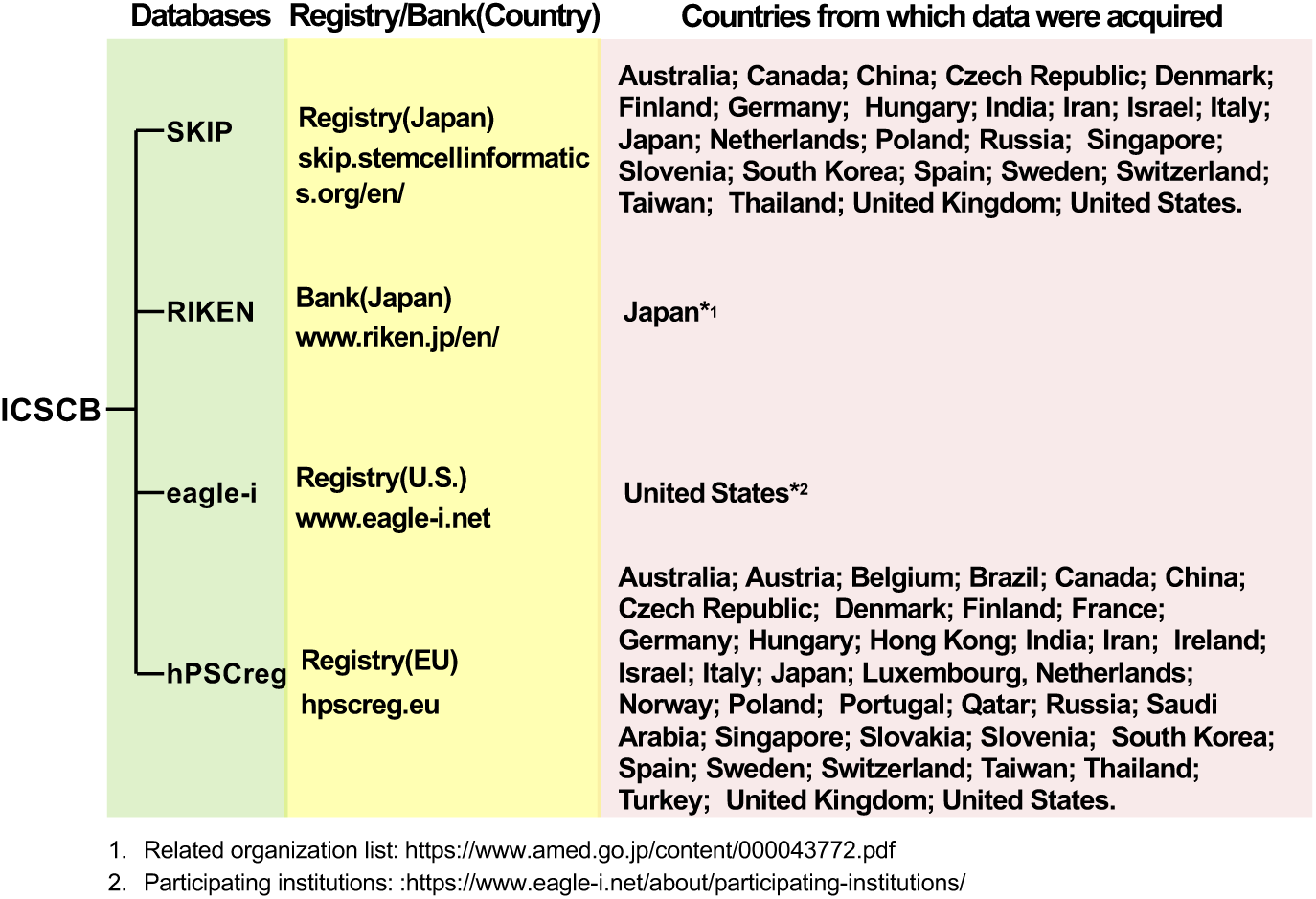
Overview of ICSCB. ICSCB includes data from three stem cell registries and one cell bank in order to maximize data coverage worldwide.

### Data integration

Since our previous research on the listed stem cell banks (Sakurai et al., 2016), the number of registered cell lines had skyrocketed from 1,483 to approximately 8,000 in the past three years. As a result, stem cell registries are tasked with collecting information on the rapidly increasing number of new cell lines and registering the cell lines into their databases as quickly as possible. However, because the stem cell banks and registries are using their own formats for data entry, the integration of the data into a centralized collection system is an extraordinary challenge. To solve this problem, we used a decentralized or distributed database system (Fujibuchi et al., 1998) by adopting items of different database formats into 16 attributes, or terms, from three MIACARM modules: stem cell general identification, donor identification, and source cell identification (**Table 2**). To practically integrate the data from the four data resources (SKIP, hPSCreg, RIKEN BRC, and eagle-i), we adopted a mechanism of cross-reference tables that allow users to conduct a search using MIACARM terms that are translated into the corresponding terms in the individual data resources to implement the search. For example, the term “Stem cell ID” in MIACARM was translated into the terms “stem cell id” (hPSCreg), “stem cell id” (SKIP), “CellID” (RIKEN BRC), and “cell line label” (eagle-i) for the search implementation. Thus, ICSCB submits search requests to each data resource with its own (translated) terms and integrates all retrieved results by common MIACARM terms, thereby achieving a standardized data format at the level of display (**Figure 4**).

**Table 2.** **Cross-reference table for integration of four databases according to MIACARM module (as of December 6, 2020)**.

| MIACARM module | ICSCB term | hPSCreg | SKIP | RIKEN BRC | eagle-i |
| --- | --- | --- | --- | --- | --- |
| Stem cell general identification | Stem cell ID | stem cell id | stem cell id | CellID | cell line label |
|  | Stem cell name | stem cell name | cell line name | CellName | cell line label |
|  | Stem cell type | N.A. | cell type | cell grouping | cell line type |
|  | Cell grade | N.A. | research grade | N.A. | N.A. |
|  | Produced by | produced by | establisher name | originator | N.A. |
|  | Provider/distributor | distributor | establisher organization | depositor | cell line provider |
|  | Reference publication | publication | pubmed ID | reference | N.A. |
| Donor identification | Gender of donor | gender | donor sex | gender | sex |
|  | Ethnicity of donor | race | donor race | race | ethnicity |
|  | Health status | health status | disease name | disease | diagnosed disease |
| Source cell identification | Source cell type | source cell type | N.A. | N.A. | N.A. |
|  | Organ/tissue of origin of source cell | origin of source cell | organ/tissue of origin of source cell | N.A. | N.A. |
| Ethical operation | Informed consent from donor | N.A. | N.A. | description | N.A. |
|  | MTA | N.A. | N.A. | description | N.A. |
| Data | Data Id | N.A. | N.A. | N.A. | N.A. |
|  | Data type | N.A. | N.A. | N.A. | N.A. |

N.A., not available.

**Fig 4:**
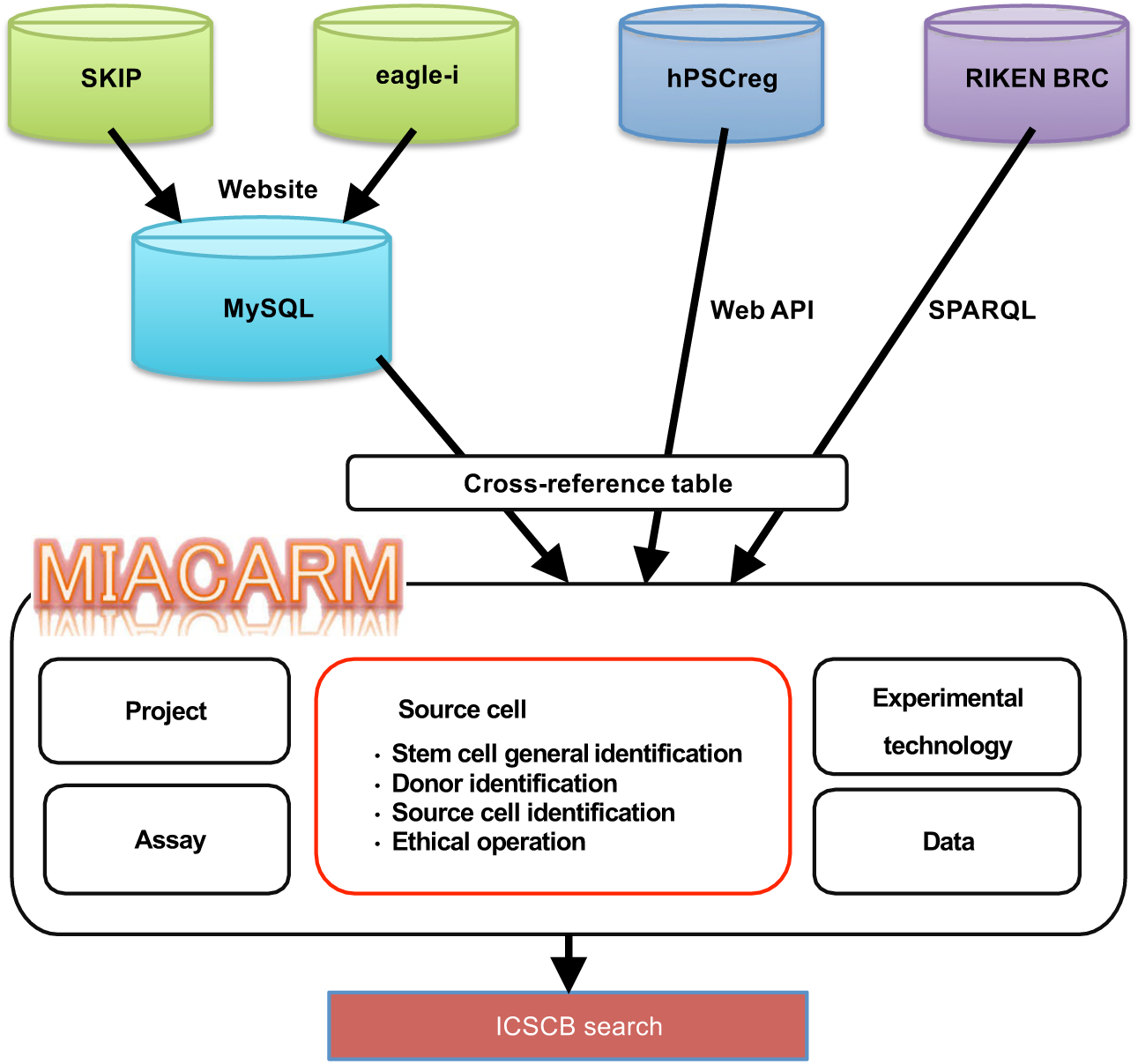
Workflow of ICSCB data integration. The SKIP and eagle-i databases were fully replicated from websites and imported to MySQL (even when updating ICSCB), whereas hPSCreg and RIKEN BRC used a web API and SPARQL for data collection, respectively. Cross-reference tables (Table 3) were used when ICSCB integrated and standardized cell line data.

### ICSCB workflow and search engine updating

In order to provide fast and easy access to the latest and accurate cell line information, we built an automatic updating system that adds newly released cell lines to ICSCB as soon as they become available in any of the four data resources. Data from eagle-i and SKIP are directly collected and stored in the MySQL database with the terms required for the MIACARM modules. Data from hPSCreg and RIKEN BRC are collected on the fly per request using a web application programming interface (API) provided by the respective sites. RIKEN BRC also uses SPARQL language for data retrieval requests (Kim et al., 2017; Kobayashi et al., 2016).

To simplify the search process, ICSCB provides an easy-to-use and mobile-friendly web application. The goal of the application is to help users find the desired stem cell lines as quickly as possible. The interface of the search engine is designed with the 16 MIACARM terms (**Table 2**) except the term “Stem cell ID”. Users receive result pages containing all the matching results listed in a table that includes all the basic attributes under the structure of MIACARM. To ensure a more specific search with a wide variety of attributes, ICSCB is designed to accommodate searches not only by standardized terms from MIACARM but also by terms specific to each of the four data resources, such as “age” or “country” (**Figure 5A**). When user queries are submitted, ICSCB simultaneously retrieves MIACARM standardized data and resource-specific data so as not to miss any relevant entries. If a keyword entered by a user in a general keyword search does not exist in MIACARM terms but is included in data specific to any of the four data resources, the user will get detailed descriptions of the matching data in the results page. For example, even if the standardized MIACARM terms do not contain “transgene”, it is still possible to enter a gene name into the keyword field (e.g., Sox2) such that the results page will display relevant entries by showing the indicated keyword in the extra field below **(Figure 5B)**. Furthermore, the user can filter the results by data resource and detailed keywords from the “Searching options” box on the results page to narrow down the results list. In addition, all results can be easily downloaded as a table directly from the results page.

**Fig 5:**
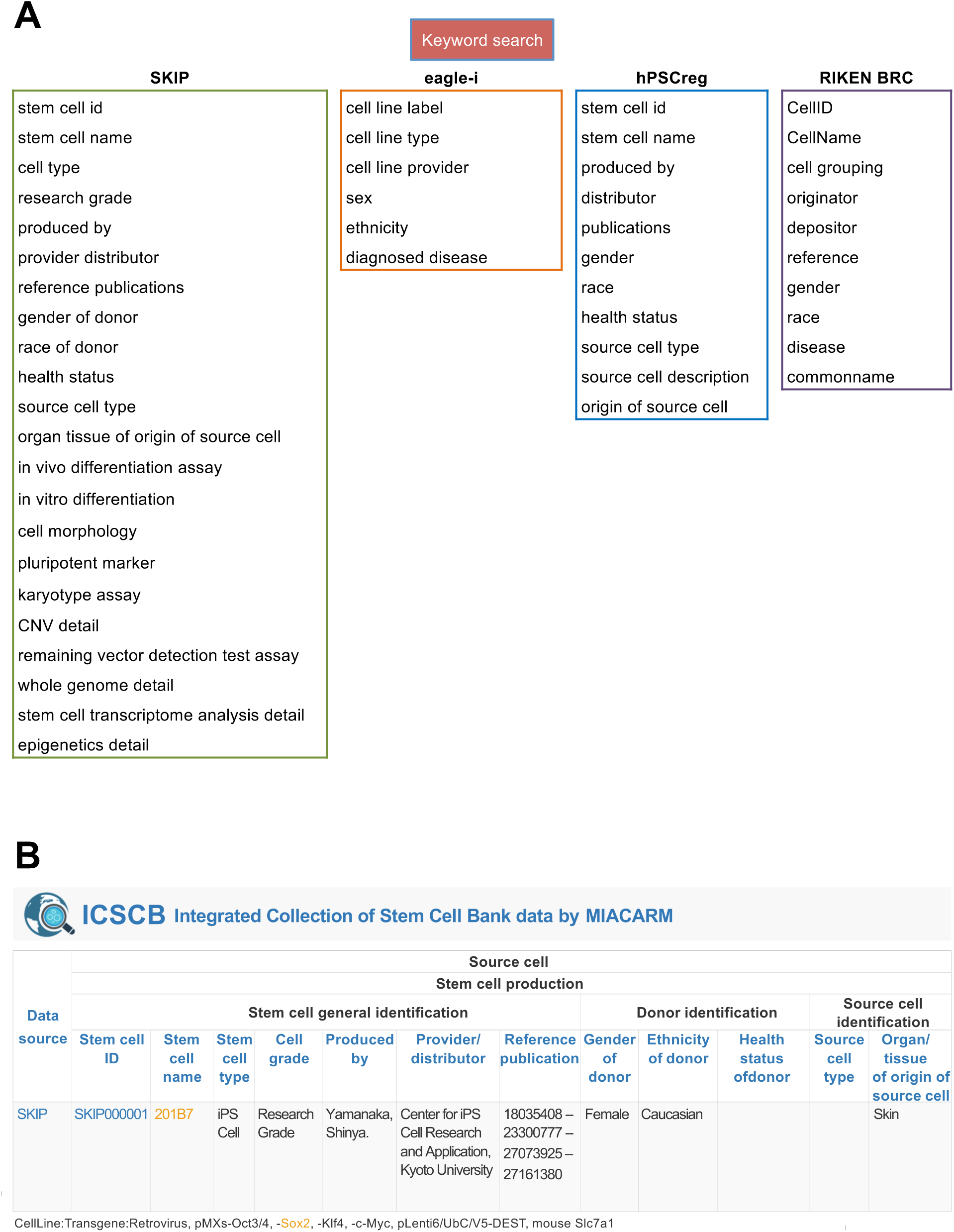
Keyword search is automatically extended to all terms provided by the four data resources even if a keyword is not included in standardized MIACARM terms. (A) Terms specific to each of the four data resources. (B) Even if the standardized MIACARM terms do not contain, for example, “transgene”, it is still possible to enter a gene name into the keyword field (e.g., Sox2), which will lead users to results from the four data resources with relevant information. The results of the match will be shown in another row below the standardized fields.

ICSCB also provides a quality control panel based on MIACARM, thereby supporting customized searches according to quality control results. At present, assays for teratoma formation, differentiation ability in vitro, morphology data, marker gene expression/surface antigen expression data, karyotyping assay results, copy number variation, residual exogene detection results, genome profiling, transcriptome profiling, and epigenome profiling data are accessible from ICSCB.

## ACKNOWLEDGEMENTS

The authors deeply appreciate Dr. Peter Karagiannis for kindly reviewing the manuscript. This work was partially supported by the Core Center for iPS Cell Research, Research Center Network for Realization of Regenerative Medicine (16bm0104001h0004) and the Formulation of Regenerative Medicine National Consortium which Renders Nation-wide Assistance to Clinical Researches, Project to Build Foundation for Promoting Clinical Research of Regenerative Medicine (19bk0204001h0004, 19bk0204001s0104), Japan Agency for Medical Research and Development (AMED); and the German Academic Exchange Service (DAAD) PPP grant.

## AUTHOR CONTRIBUTIONS

YC and YP drafted the manuscript. SM, TM, HO, JD, SS, AK, HM, YN, MS, JS, and WF provided and facilitated the stem cell data. WF and KS conceptualized the research. AK, GS, and WF led the project.

## COMPETING INTERESTS

HO is a founding scientist of SanBio Co., Ltd. and K Pharma Inc.

## Notes

### Competing Interest Statement

We have one potential conflict of interest to disclose: Professor Hideyuki Okano of Keio University, Japan, is a founding scientist of SanBio Co. Ltd. and K Pharma Inc.

### Summary of Updates

Concluding remarks and future plan in Results and Discussion updated

