## Supplementary Information for "Integrated Collection of Stem Cell Bank data, a data portal for standardized stem cell information"

### SUPPLEMENTAL FIGURES

**Fig S1. Related to Figure 2.**

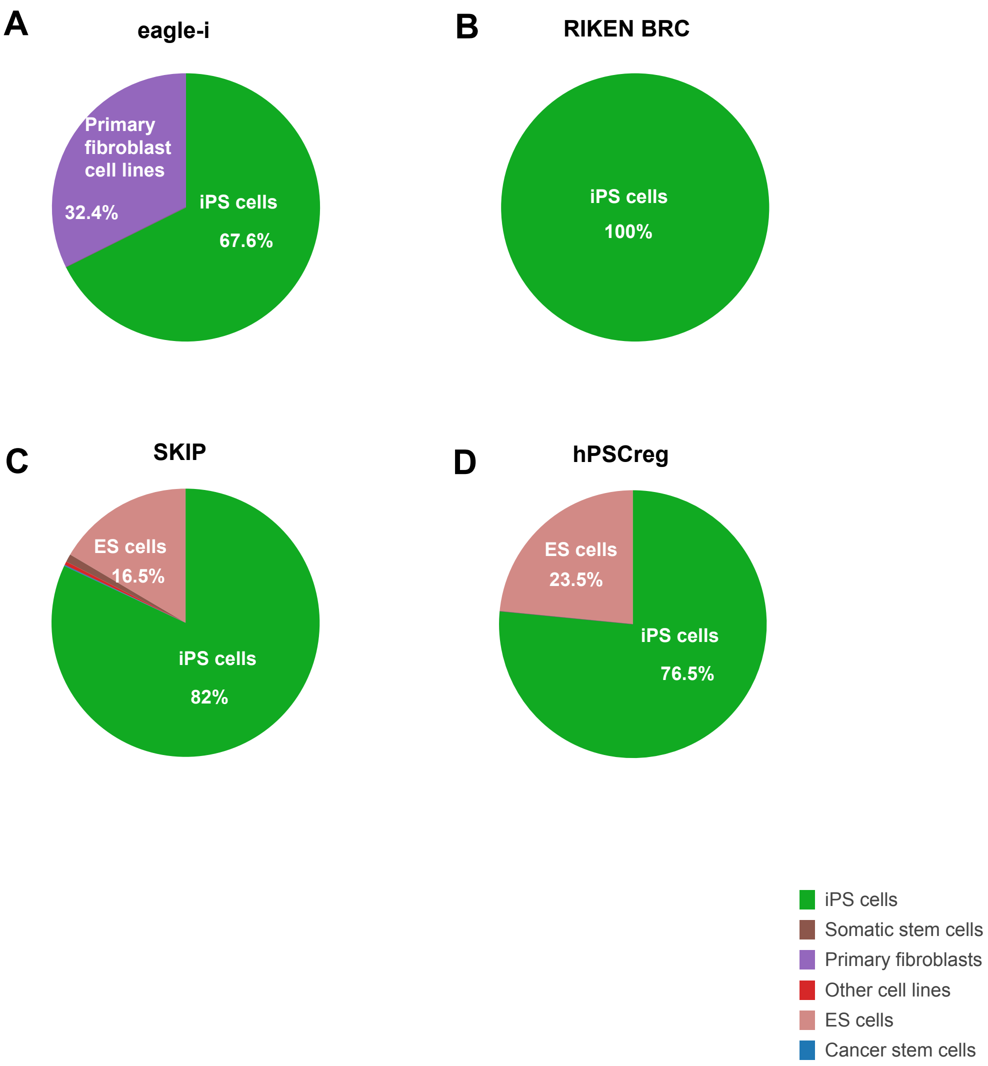


**Fig S1. Details of cell line types collected by eagle-i, RIKEN BRC, SKIP, and hPSCreg** (as of December 6, 2020)**.** (A) eagle-i, (B) RIKEN BRC, (C) SKIP, and (D) hPSCreg.

**Fig S2. Related to Figure 2.**

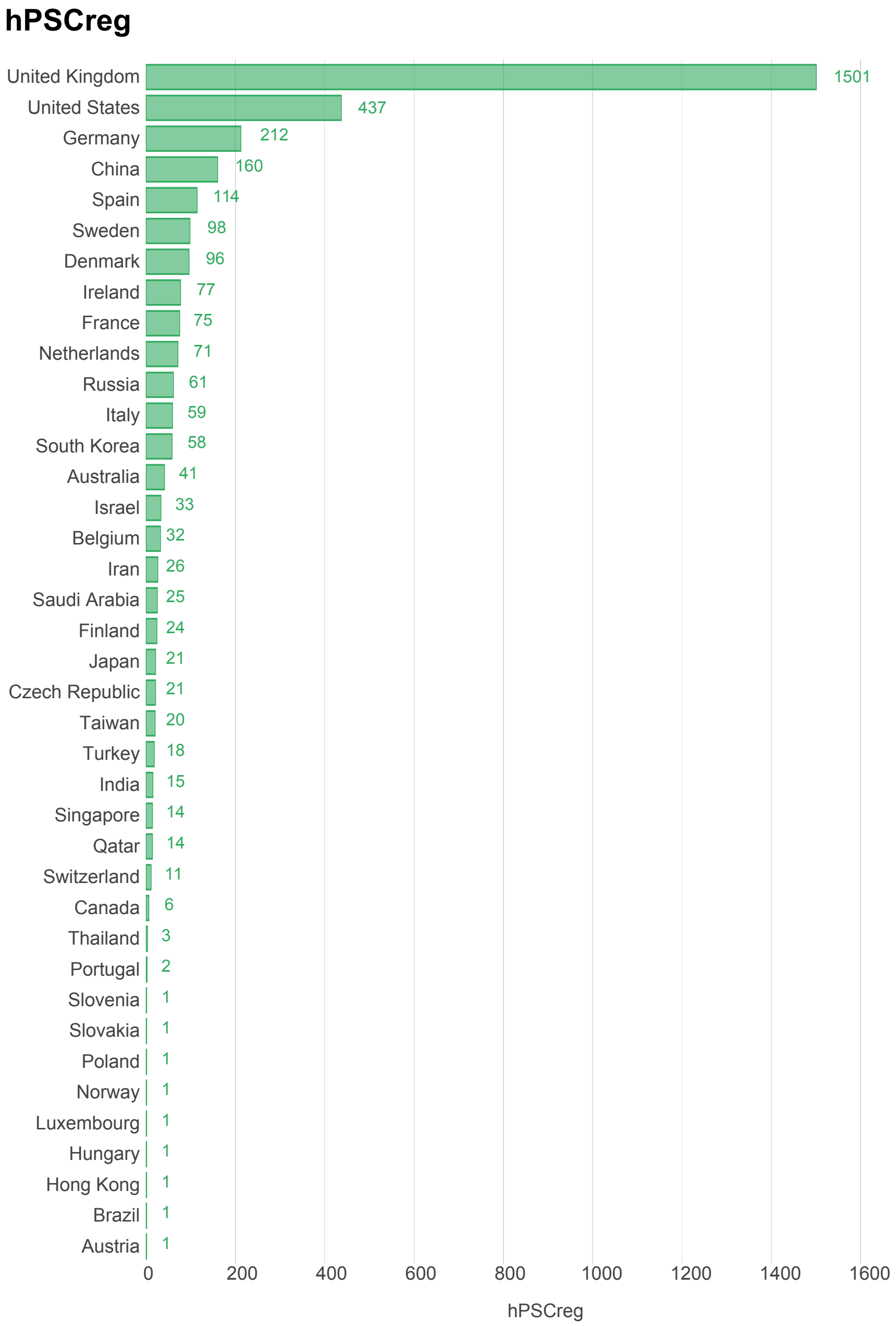


**Fig S2. Details of countries that have established cell lines in hPSCreg** (as of December 6, 2020)**.**

**Fig S3. Related to Figure 2.**

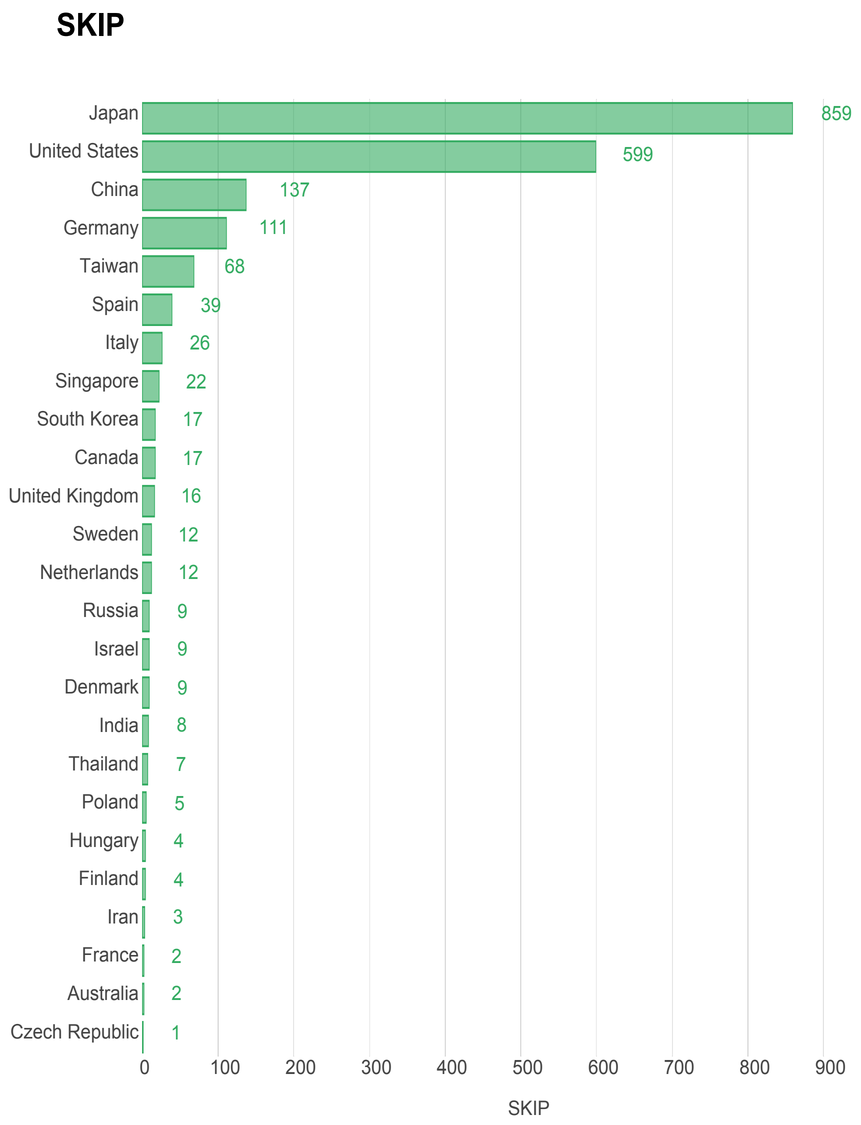


**Fig S3. Details of countries that have established cell lines in SKIP** (as of December 6, 2020)**.**

### SUPPLEMENTAL TABLES

#### Table S2. Related to Figure 2A and Figure S1.

| Stem_cell_type | hPSCreg | SKIP | RIKEN BRC |
|---|---|---|---|
| **ES cells** | 788 | 927 | 0 |
| **iPS cells** | 2572 | 4603 | 3548 |
| **Somatic stem cells** | 0 | 54 | 0 |
| **Cancer stem cells** | 0 | 7 | 0 |
| **Primary fibroblast cell lines** | 0 | 0 | 0 |
| **Others** | 0 | 24 | 0 |
| **Total** | **3360** | **5615** | **3548** |

**Table S2. List of cell line types across all four databases** (as of December 6, 2020)**.**

**Table S3. Related to Figure 2B.**

| Health_status | Count |
|---|---|
| **Healthy** | 9708 |
| **Diseased** | 6363 |
| **Total** | **16071** |

**Table S3. Statistics of healthy/diseased cell lines in ICSCB** (as of December 6, 2020)**.**

### Table S4. Related to Figure 2C.

| Country | SKIP | hPSCreg | Total |
|---|---|---|---|
| United Kingdom | 16 | 1501 | 1517 |
| United States | 599 | 437 | 1036 |
| Japan | 859 | 21 | 880 |
| Germany | 111 | 212 | 323 |
| China | 137 | 160 | 297 |
| Spain | 39 | 114 | 153 |
| Sweden | 12 | 98 | 110 |
| Denmark | 9 | 96 | 105 |
| Taiwan | 68 | 20 | 88 |
| Italy | 26 | 59 | 85 |
| Ireland | 0 | 77 | 77 |
| South Korea | 17 | 58 | 75 |
| Netherlands | 12 | 71 | 83 |
| Russia | 9 | 61 | 70 |
| France | 2 | 75 | 77 |
| Australia | 2 | 41 | 43 |
| Israel | 9 | 33 | 42 |
| Singapore | 22 | 14 | 36 |
| Belgium | 0 | 32 | 32 |
| Iran | 3 | 26 | 29 |
| Canada | 17 | 6 | 23 |
| Finland | 4 | 24 | 28 |
| Czech Republic | 1 | 21 | 22 |
| India | 8 | 15 | 23 |
| Turkey | 0 | 18 | 18 |
| Qatar | 0 | 14 | 14 |
| Switzerland | 0 | 11 | 11 |
| Thailand | 7 | 3 | 10 |
| Hungary | 4 | 1 | 5 |
| Poland | 5 | 1 | 6 |
| Portugal | 0 | 2 | 2 |
| Saudi Arabia | 0 | 25 | 25 |
| Austria | 0 | 1 | 1 |
| Brazil | 0 | 1 | 1 |
| Luxembourg | 0 | 1 | 1 |
| Norway | 0 | 1 | 1 |
| Slovakia | 0 | 1 | 1 |
| Slovenia | 0 | 1 | 1 |
| Hong Kong | 0 | 1 | 1 |
| **Total** | **1998*** | **3354**** | **5352** |

*The number was calculated by 5615 (total) - 3647 (hPSCreg or eagle-i or unknown country) + 30 (dual country) = 1998.

**The number was calculated by 3360 (total) - 6 (synonyms) = 3354

**Table S4. Statistics of cell line types based on country** (as of December 6, 2020)**.**

**Table S5. Related to Figure 2D.**

| Disease Category in MeSH | Count of diseases |
|---|---|
| **Urogenital Diseases** | 26 |
| **Skin and Connective Tissue Diseases** | 12 |
| **Others** | 16 |
| **Nutritional and Metabolic Diseases** | 21 |
| **Nervous System Diseases** | 614 |
| **Neoplasms** | 16 |
| **Musculoskeletal Diseases** | 52 |
| **Mental Disorders** | 90 |
| **Male Urogenital Diseases** | 25 |
| **Immune System Diseases** | 15 |
| **Hemic and Lymphatic Diseases** | 26 |
| **Genetic Disorders** | 179 |
| **Eye Diseases** | 52 |
| **Endocrine System Diseases** | 108 |
| **Digestive System Diseases** | 3 |
| **Chemically Induced Disorders** | 4 |
| **Cardiovascular Diseases** | 107 |
| **Total** | **1366** |

**Table S5. Statistics of diseased cell lines based on disease category** (as of December 6, 2020)**.**

### EXPERIMENTAL PROCEDURES

#### Generation of Fig. 1

Among all the databases, SKIP and hPSCreg provided details of countries from which data were acquired. For SKIP, this information was provided on its homepage (skip.stemcellinformatics.org/en/). For hPSCreg, country information for every cell line could be accessed from its homepage (https://hpscreg.eu/) by clicking “find by location”.

#### Generation of Fig. 2

Full information data were directly downloaded from ICSCB results page (Table S1) and filtered according to the following criteria: (A) stem cell type (Table S2); (B) health/disease status (Table S3); (C) country (Table S4); and (D) disease (Table S5). Disease categories were determined by search results with keywords under the “Disease Category” in NCBI MeSH page (https://www.ncbi.nlm.nih.gov/mesh). For example, searching with keywords of “Parkinson disease” will lead to the MeSH term “Nervous System Diseases” under the “Disease Category”. Pie charts and bar graphs were produced by R (graph.r) using the package “plotly”.
